# The neurons that drive infradian sleep-wake and mania-like behavioral rhythms

**DOI:** 10.1101/2023.11.14.566955

**Authors:** Pratap S. Markam, Clément Bourguignon, Lei Zhu, Martin Darvas, Paul V. Sabatini, Maia V. Kokoeva, Bruno Giros, Kai-Florian Storch

## Abstract

Infradian mood and sleep-wake rhythms with periods of 48 hr and beyond have been observed in bipolar disorder (BD) subjects that even persist in time isolation, indicating an endogenous origin. Here we show that mice exposed to methamphetamine (Meth) in drinking water develop infradian locomotor rhythms with periods of 48 hr and beyond which extend to sleep length and mania-like behaviors in support of a model for cycling in BD. This cycling capacity is abrogated upon genetic disruption of DA production in DA neurons of the ventral tegmental area (VTA) or ablation of nucleus accumbens (NAc) projecting, dopamine (DA) neurons. Chemogenetic activation of NAc-projecting DA neurons leads to locomotor period lengthening in clock deficient mice, while cytosolic calcium in DA processes of the NAc was found fluctuating synchronously with locomotor behavior. Together, our findings argue that BD cycling relies on infradian rhythm generation that depends on NAc-projecting DA neurons.

## Introduction

Rhythms in sleep-wake are typically aligned with the solar day, adhering to a periodicity of 24 hr. However, there are also accounts of sleep-wake cycling with periods longer than a day. Individuals affected by “non–24-hour sleep–wake rhythm disorder” (N24SWD) ^1^, exhibit sleep-wake rhythms that can reach far into the infradian (>>24 hr) period range ^2^. Interestingly, infradian sleep-wake cycles with the particular period of 48 hr have been specifically reported in the context of bipolar disorder (BD) ^3 4,5^. Here sleep length varies between successive days and this sleep cycling typically goes hand in hand with mood states alternating between (hypo)mania on short-sleep and depression on long-sleep days. BD 48 hr cycling was shown to persist even under conditions of time isolation ^4^, indicating an endogenous origin, however, as with infradian N24SWD, the biological basis of BD-associated 48 hr cycling is unknown.

Sleep-wake rhythms longer than 24 hr can be experimentally induced in laboratory animals upon chronic treatment with methamphetamine (Meth) via drinking water ^6,7^. Upon Meth exposure, a second rhythmic component (2ndC) emerges along-side the circadian rhythm, which typically develops over the course of several days to weeks and whose period is affected by the concentration of Meth. Such 2ndC can be produced even in the absence of the canonical circadian timer underscoring its independence from the clock machinery ^8 9^. A 2ndC can also emerge upon constant Meth infusion via implanted mini pumps ^8^, arguing against rhythmic Meth uptake through intermittent drinking as a driver of the 2ndC. A key target of Meth is the dopamine transporter (DAT), which is set into inverse action upon Meth exposure, turning it into a DA releasing entity, thereby elevating extracellular DA ^10^. Consistently, *DAT* gene disruption, which also causes an increase in extracellular DA^11^, equally leads to the emergence of a 2ndC ^12^, a finding which also demonstrates that 2ndC production does not solely rely on Meth treatment. Together, these data suggest that DA neurons, which are the near exclusive sites of DAT expression, exert a crucial role in the genesis of a second rhythmic locomotor component ^12^, possibly harboring a dopamine-based oscillator (DO) that drives it. Here we set out to identify the neuronal substrate of the oscillator process and to find evidence in mice for a role of the oscillator in driving mood and sleep cycling in BD.

## Results

### Infradian rhythm emergence by chronic methamphetamine treatment

In the past, Meth-induced 2ndCs were generally viewed as additional circadian rhythms as they typically exhibited periods not far from 24 hr. However, most of these studies used Meth at concentrations up to 50mg/L drinking water, which may account for the rather limited period range that was observed^13^. Here we examined running wheel activity of mice exposed to increasing levels of Meth, up to 100mg/L, which is twice the concentration normally used (Figure 1A). As expected, Meth treatment led to the emergence of a 2ndC over time as demonstrated by the pattern of running wheel activity and associated periodogram analysis (Figure 1B). However, time of 2ndC emergence with respect to Meth treatment onset was found variable, with some animals readily showing a 2ndC, while others took weeks to respond (Figure 1C). There was also considerable variability in the period the 2ndC adopts in response to Meth, with faster responding animals leaning towards longer period 2ndCs (Figure 1C). Generally, 2ndC emergence is preceded by a gradual lengthening of the main (circadian) locomotor component (Figure 1B, dashed white box in CWT displays right next to actograms). Confirming previous findings ^14-16^, the 2ndC can gradually or abruptly change its period over time (Figures 1B and 1C) which can be viewed as reflections of a rather labile oscillator that drives the 2ndC. Figure 1B (right) further demonstrates another known feature of the second component: relative coordination with the circadian timer, i.e., the period of the 2ndC changes depending on its phase angle position relative to the circadian timer ^6,7^. Importantly, upon prolonged administration of 100mg/L Meth, we found a high fraction of animals to exhibit 2ndCs with periods in the infradian range (Figures 1C-1F), with a considerable portion (≥30%) reaching periods beyond 40 hr (Figures 1E). The relatively even distribution of infradian peaks across a wide infradian range demonstrated by the heatmap display of individual periodograms from animals under Meth treatment (Figure 1D) underscores a profound period-flexibility. This is reflective of a highly tunable rhythms generator^12^, which is in stark contrast to the circadian oscillator whose period range is rather constrained to near 24 hr periodicities. Figure 1F shows actograms plotted in modulo according the ultimate period the animals adopt, illustrating the regularity and robustness of Meth-induced infradian sleep-wake rhythms. Taken together, we show that chronic Meth treatment leads to the emergence of a rhythmic locomotor component of varying periodicities that often reaches far into the infradian range, arguing that the oscillator that drives the 2ndC is not circadian in nature.

**Figure 1.**
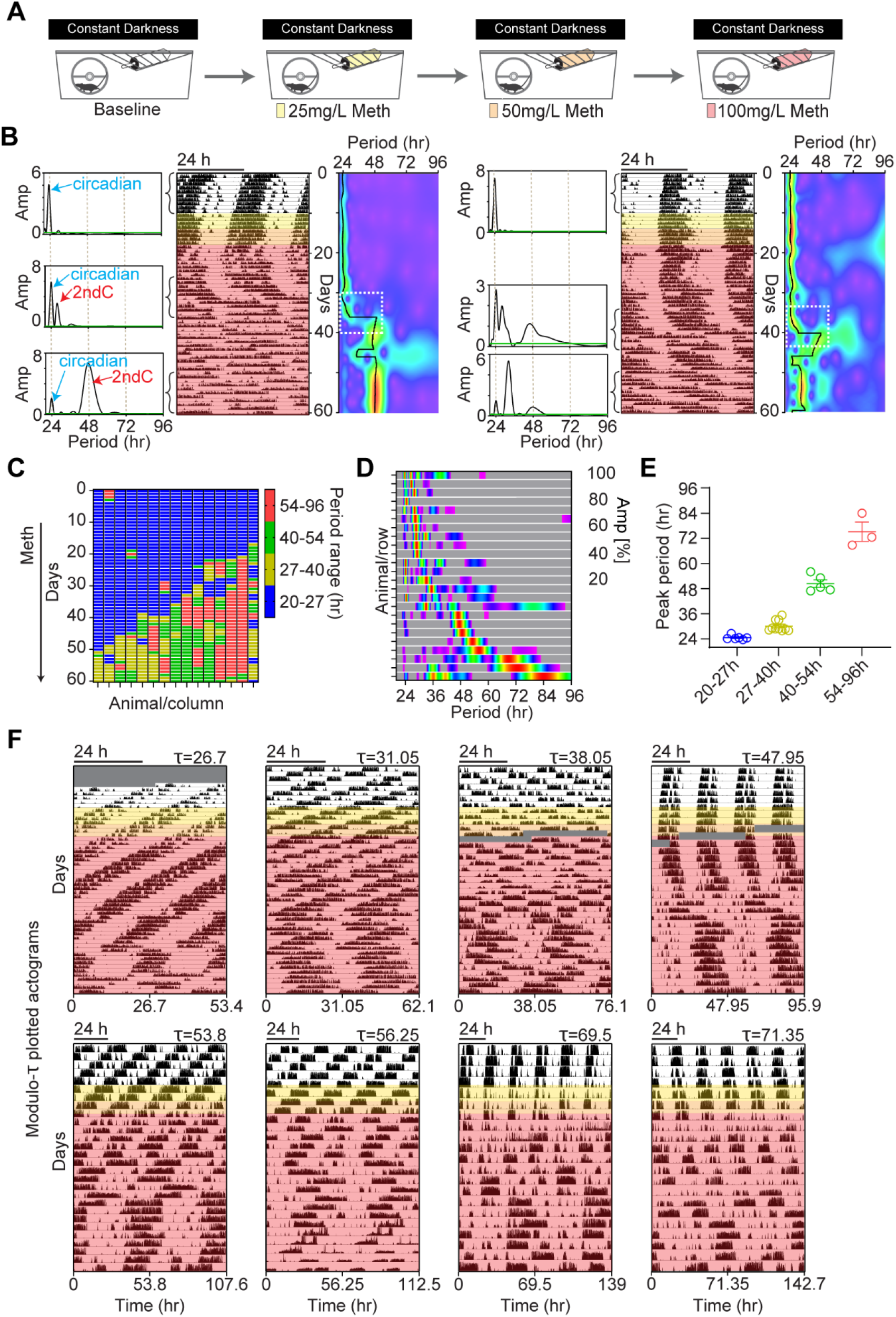
Variability and extent of infradian rhythm induction by methamphetamine. (A) Experimental regimen of running wheel activity monitoring in response to escalating concentrations of Meth in drinking water. (B) Representative actograms displaying running wheel activity patterns of two single housed animals exposed to Meth. Meth concentrations applied are indicated by color shading in correspondence to A). Lomb-Scargle (LS) periodograms on the left and continuous wavelet transform (CWT) heatmaps on the right of each actogram show the emergence and evolution of infradian locomotor rhythms. Green line in periodograms demarcates the significant threshold for rhythmicity. Black traces in heatmaps are the wavelet ridges reporting the instantaneous peak period. (C) Evolution of the peak period across animals identified by the wavelet ridge of the CWT. Time course of Meth treatment as indicated in B. (D) Composite heatmap of normalized periodograms from the final 14 days of Meth treatment suggests profound period flexibility. (E) Binned display of highest peaks derived from periodograms shown in D. Shown are means ± SEM. (F) Representative actograms of Meth-treated mice plotted modulo according the peak period from the last 14 days of the 60-day actogram. The modulo actograms demonstrate that mice can adopt robust rhythmicity in the near to far infradian range upon Meth exposure via drinking water. Greyed areas indicated data loss.

### Infradian rhythmic mice as model for bipolar cycling

To demonstrate that similar to 48 hr cycling BD patients^3-5^, also the 48 hr rhythmicity in mice extends to sleep, we examined the effect of chronic Meth treatment on spontaneous locomotor activity recorded by passive infrared sensors (PIRs) (Figure 2A). Using this approach, we found that even in the absence of a running wheel, mice consistently adopted 48 hr rhythmicity in locomotor activity in response to Meth (Figures 2B and S1A). We then transformed the PIR data into sleep scores in accordance to a procedure that has been validated against EEG-derived sleep ^17,18^. The resulting sleep pattern (Figure 2B, right) showed a 48 hr rhythmicity expectedly mirroring the PIR-derived activity pattern (Figure 2B, left). Figure 2C confirms that infradian spectral power in sleep rhythmicity is virtually absent prior to Meth exposure but profoundly present when mice exhibited 48 hr cycling (see also Figure S1B and S1C). When comparing successive days during 48 hr cycling, we found sleep length to differ strongly. On active days, mice spent 30-40% of the day sleeping, whereas they allocated 60-80% of the day to sleep on inactive days or days prior to Meth treatment (Figure 2D), which agrees well with observations in 48 hr cycling bipolar subjects who alternate between short sleep and a rather standard sleep length ^4,5^. Consistently, total sleep during 48hr cycling was lower when compared to pre-Meth conditions (Figure 2E), in support of a substantially reduced sleep need in 48 hr cycling mice or humans.

**Figure 2.**
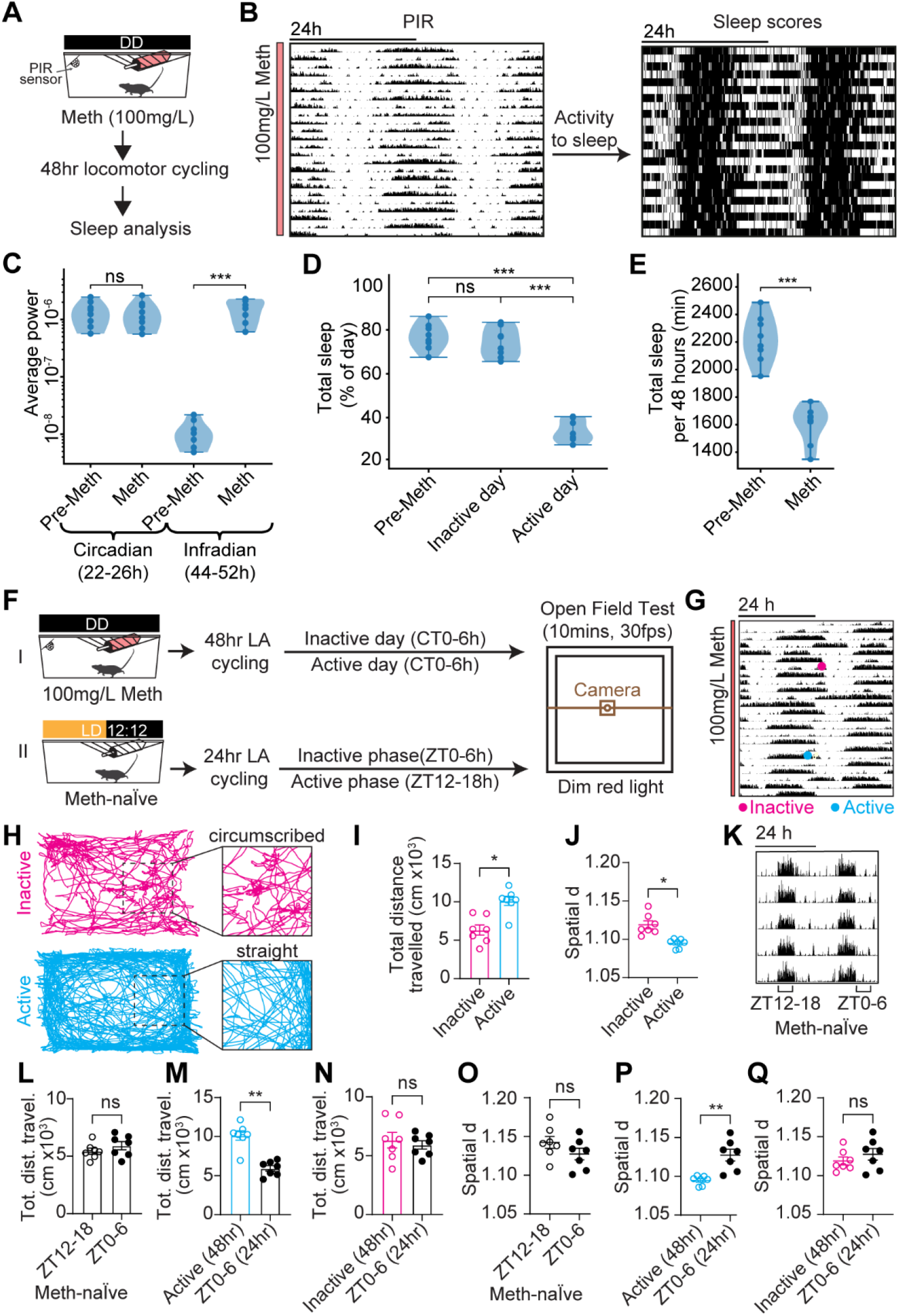
48 hr rhythmic mice show cycling in sleep and mania-associated behaviors. (A) Regimen to record sleep during 48 hr rhythmicity. (B) PIR-derived spontaneous locomotor activity (left) and sleep (right) of a 48 hr rhythmic mouse. (C) Rhythmic strength (Scale-averaged spectral power) in the circadian (22-26 hr) and infradian (44-52 hr) range prior to Meth exposure and during 48 hr cycling. (D) Daily total sleep prior to Meth and on active and inactive days while cycling. (E) Daily sleep (average across four consecutive days) prior to Meth and during 48 hr cycling. (F) Experimental design for open field test (OFT) in 48 hr and 24 hr-rhythmic mice. (G, H) Actogram (G) of a 48 hr cycling mouse used for OFT analysis. Time of testing during active/inactive days indicated. (H) Representative locomotor traces derived from 10 min open field video recording on inactive and active days, as indicated in (G). (I) Total distance travelled during OFT of 48 hr cycling mice during their active and inactive days. (J) Spatial d in 48 hr cycling mice during inactive and active days. (K) Representative actogram of Meth-naïve animals displaying 24hr rhythms in locomotor activity. (L) Total distance travelled during OFT conducted during the active (ZT12-18) and inactive (ZT0-6) phases of 24 hour rhythmic, Meth-naïve mice. (M, N) Total distance travelled on active (M) and inactive (N) days by 48 hr cyclers compared to the inactive phase of Meth-naïve mice. (O) Spatial d in Meth-naïve mice during their active vs inactive phase. (P, Q) Spatial d on active (P) and inactive (Q) days by 48 hr cyclers compared to the inactive phase of Meth-naïve mice. Mean ± SEM. n= 8 (C-E) and 7 (I, J, L-Q). Mann-Whitney test, C-E and L-Q. Wilcoxon’s test, I and J. ns, not significant. \**P*<0.05; **P<0.01; \*\*\**P*<0.001. See also Figure S1

To further corroborate the validity of 48 hr rhythmic mice as model for 48 hr cycling in BD, we assessed if these mice show evidence of hyperactivity every other day using the open field test. As hyperactivity or psychomotor agitation is a hallmark of BD mania ^19^, such finding would be consistent with these mice entering a mania-like state every other day. Indeed, open field video-tracking revealed that distance traveled over time was elevated when 48 hr cycling animals were tested on days when they slept little (active days) compared to the alternating days of long sleep (inactive days) (Figures 2F-I). In contrast, no difference was found in Meth-naïve animals, when tested during their corresponding active (ZT12-18) and inactive (ZT0-6) phases in line with previous findings^20^ (Figures 2K and 2L). Notably, distance traveled on active days in 48 hr cyclers was also greater when compared to Meth-naïve mice (Figure 2M), while there was no difference between inactive days and Meth-naïve animals (Figure 2N). We next examined the degree of straightness of the movement path in the open field arena. It has been previously shown that manic BD subjects would exhibit more straight-path movements over control subjects when their locomotion was traced via a room-ceiling installed video camera ^21,22^. Path straightness was quantified using the metric spatial d, which reports on the difference of spatial displacement in 2D space for a given distance traveled. The more straight a movement path, the lower spatial d and consequently spatial d was found to be lower in manic patients over healthy controls^21,22^. We applied the spatial d analysis to the open field recordings of the 48 hr rhythmic mice and found that spatial d was lower on active days when compared to inactive days (Figure 2J). This result is consistent with the visual inspection of the movement path, showing more circumscribed locomotion on inactive days when compared to active days (Figure 2H). As with distance traveled (Figure 2L), there were no differences in spatial d in Meth-naïve mice between their active and inactive phases (Figure 2O) and there was no significant difference between spatial d measured during inactive days in 48 hr cyclers vs Meth-naïve mice (Figure 2Q). However, 48hr cyclers on active days showed significantly lower spatial d as compared to Meth-naïve mice (Figure 2P). Thus, 48 hr rhythmic mice show hyperactivity and more straight path ambulation every other day, which is in line with the behavior of patients in mania, corroborating 48 hr rhythmic mice as a suitable model for mania cycling in BD.

### ^VTA^DA neurons are necessary for infradian rhythm emergence

We showed previously that chemogenetic activation of midbrain DA neurons results in period lengthening in clock-deficient mice under constant darkness conditions ^12^. Furthermore, optogenetic activation of TH-expressing VTA neurons promotes wakefulness even shortly after lights on, when daily sleep pressure is highest in mice ^23^, which suggests that ^VTA^DA neurons can override circadian sleep-wake control. Together, these findings are consistent with ^VTA^DA neurons acting as drivers of infradian rhythms in sleep-wake and mood.

To test this, we first aimed to genetically ablate ^VTA^DA neurons by employing a virus that Cre-dependently expresses genetically engineered Caspase 3 (AAV-flex-taCasp3-TEVp)^24^ to trigger cell autonomous apoptosis. To limit cell ablation to DA neurons, we injected the virus bilaterally into the VTA of *DAT-Cre* mice ^25^ (Figure 3A). Subsequent immuno-staining against the catecholamine cell marker tyrosine hydroxylase (TH) revealed a preferential loss of TH in the VTA, but not in the adjacent substantia nigra (SN) (Figures 3B and 3C), indicative of successful ^VTA^DA neuron ablation. Notably, TH expression was found retained in some cells of the ventro-medial VTA in ^VTA^Casp3 mice (Figure 3C), suggestive of poor Cre activity which we confirmed when crossing *DAT-Cre* mice with a reporter line (Figures S2A and S2B) or to *TH^fl/fl^* mice (Figure S2C). This recombination deficit also resonates with previous observations of lower DAT expression in TH cells of this ventral-medial VTA region ^26^. Next, we monitored running wheel activity of ablated animals (^VTA^Casp3 mice) in response to Meth in constant darkness (DD) to prevent interference of endogenous rhythm generation by the light-dark cycle. A gradual increase of Meth in drinking water led to the expected emergence of an infradian component in control animals, as evidenced by the distinct pattern changes in locomotor activity (Figure 3D) and corresponding emergence of >24 hr peaks in the periodogram (shown underneath actogram for the time span indicated by vertical blue bar). By contrast, ^VTA^Casp3 mice showed a profound loss of 2ndC induction capacity (Figure 3E), confirmed by the lack of a rhythmicity shift into the infradian range (Figure 3F). This was quantified by calculating the amplitude spectral density (ASD) for the circadian (20-27hr) and infradian (27-96hr) ranges, respectively, demonstrating a lack of a shift of rhythmic power into the infradian range in ^VTA^Casp3 mice upon Meth treatment (Figures 3G and 3H). Consistently, the highest periodogram peak shifted to infradian periods in controls, while it remained circadian in ^VTA^Casp3 animals (Figure 3J). Notably, total activity tended to be lower in ^VTA^Casp3 mice before and during Meth treatment (Figure 3K), suggesting an additional global effect on locomotor activity upon DA neuron ablation in the VTA, in keeping with their role in arousal maintenance ^23^. However, DA neuron ablation in the VTA did not affect circadian period based on the highest peak analysis pre-Meth (Figure 3I), indicating that the central circadian pacemaker residing in the suprachiasmatic nucleus (SCN) remained functionally intact.

**Figure 3.**
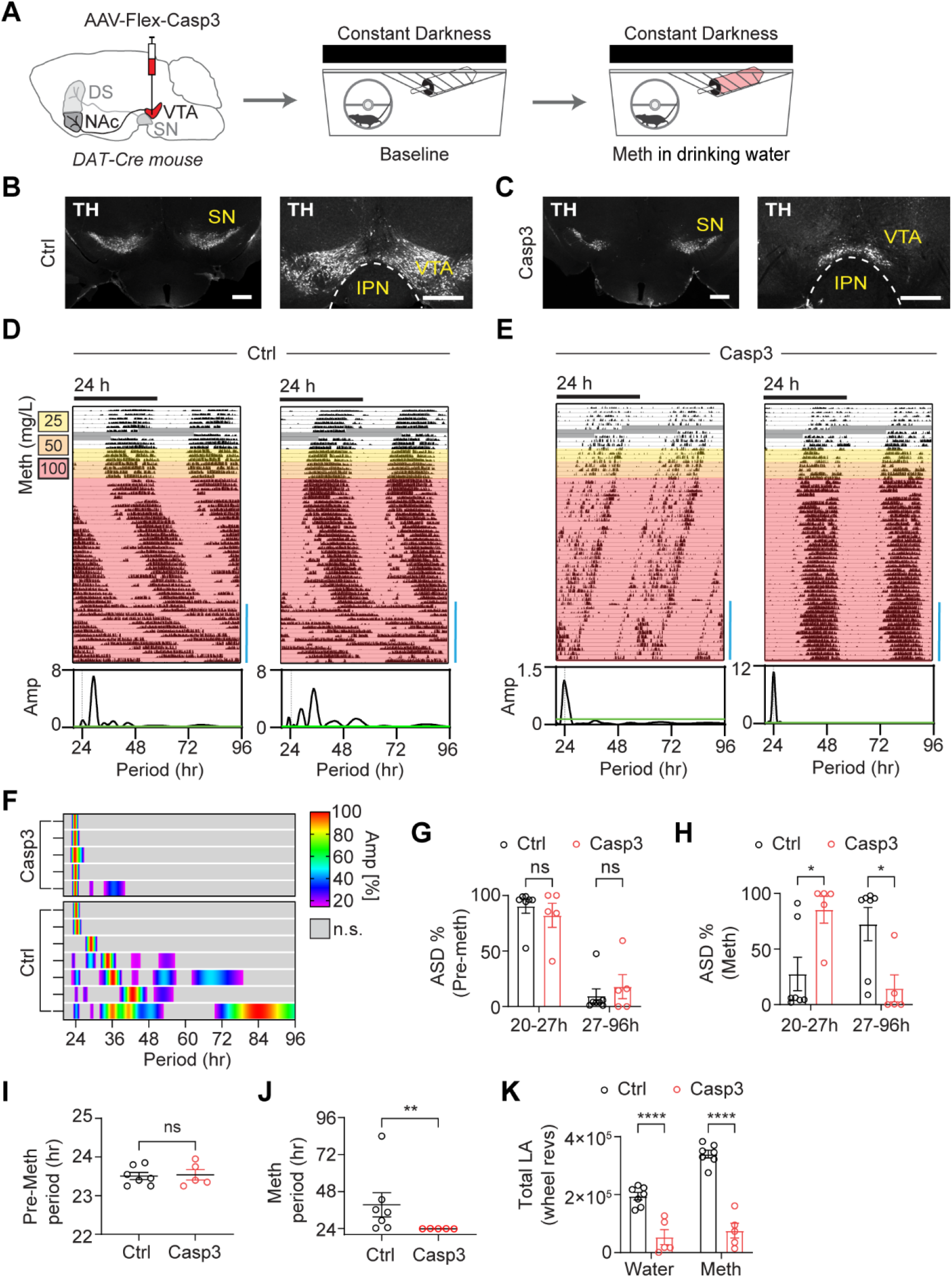
Infradian rhythm generation capacity is abrogated upon ^VTA^DA neuron ablation. (A) Experimental design: DA neuron ablation in the VTA via Casp3 and subsequent behavioral monitoring using running wheels. (B, C) Bilateral injection of AAV-DIO-taCasp3-TEVp into the VTA of *DAT-Cre* mice led to a loss of TH immunofluorescence in the VTA (C) compared to saline injected mice (B). IPN, interpeduncular nucleus; SN, substantia nigra compacta. Scale bars, 500 μm. (D, E) Representative actograms showing running wheel activity of ^VTA^Casp3 mice and controls in constant darkness in response to escalating levels of Meth in drinking water. LS-periodograms below the actograms are computed for the time window indicated by the blue bar at the bottom right of the actograms. Greyed areas indicate data loss. (F) Composite display of periodograms from individual animals computed from the final 2 weeks of recording under Meth treatment. Periodograms are normalized to its peak amplitude. Grey shading indicates absence of significant amplitudes. (G, H) Periodogram-derived amplitude spectral density (ASD) of significant periodicities in the circadian (20-27 hr) and infradian (27-96 hr) period range before (G) and during the final 2 weeks of Meth-exposure (H). (I, J) Periodogram-derived highest peak in the 20-96 hr range before (I) and during the final 2 weeks of Meth-treatment (J). (K) Total locomotor activity (wheel revolutions) prior to (Water) and during Meth treatment (Meth, final 2 weeks of treatment), Mean ± SEM. n= 5-7. Mann-Whitney’s test, I and J. Two-way ANOVA with Bonferroni’s multiple comparison test, G, H and K. ns, not significant; *P<0.05; **P<0.01; ****P<0.0001. See also Figure S2.

### ^VTA^TH is required for infradian rhythmicity

Considering that Meth-treatment as well as DAT-deficiency alters DA levels ^11^ (Figure 4A), it seemed likely that DA would be of critical importance for 2ndC induction. We therefore disrupted tyrosine hydroxylase (TH), an enzyme essential for DA biosynthesis ^27^, exclusively in ^VTA^DA neurons (Figure 4B). Adult *Th^fl/fl^ mice* ^28^ were injected with a Cre recombinase expressing AAV (AAV-GFP-Cre) into the VTA bilaterally, which led to a selective loss of TH in the VTA (Figures 4C, 4D, and S3). As in the case of ^VTA^Casp3 mice, ^VTA^THKO mice also failed to produce a 2ndC in response to Meth treatment (Figures 4E-4K), while the circadian period pre-Meth remained unaffected (Figure 4J). Together, these results indicate that DA neurons of the VTA and more specifically their DA production capacity is required for 2ndC emergence.

**Figure 4.**
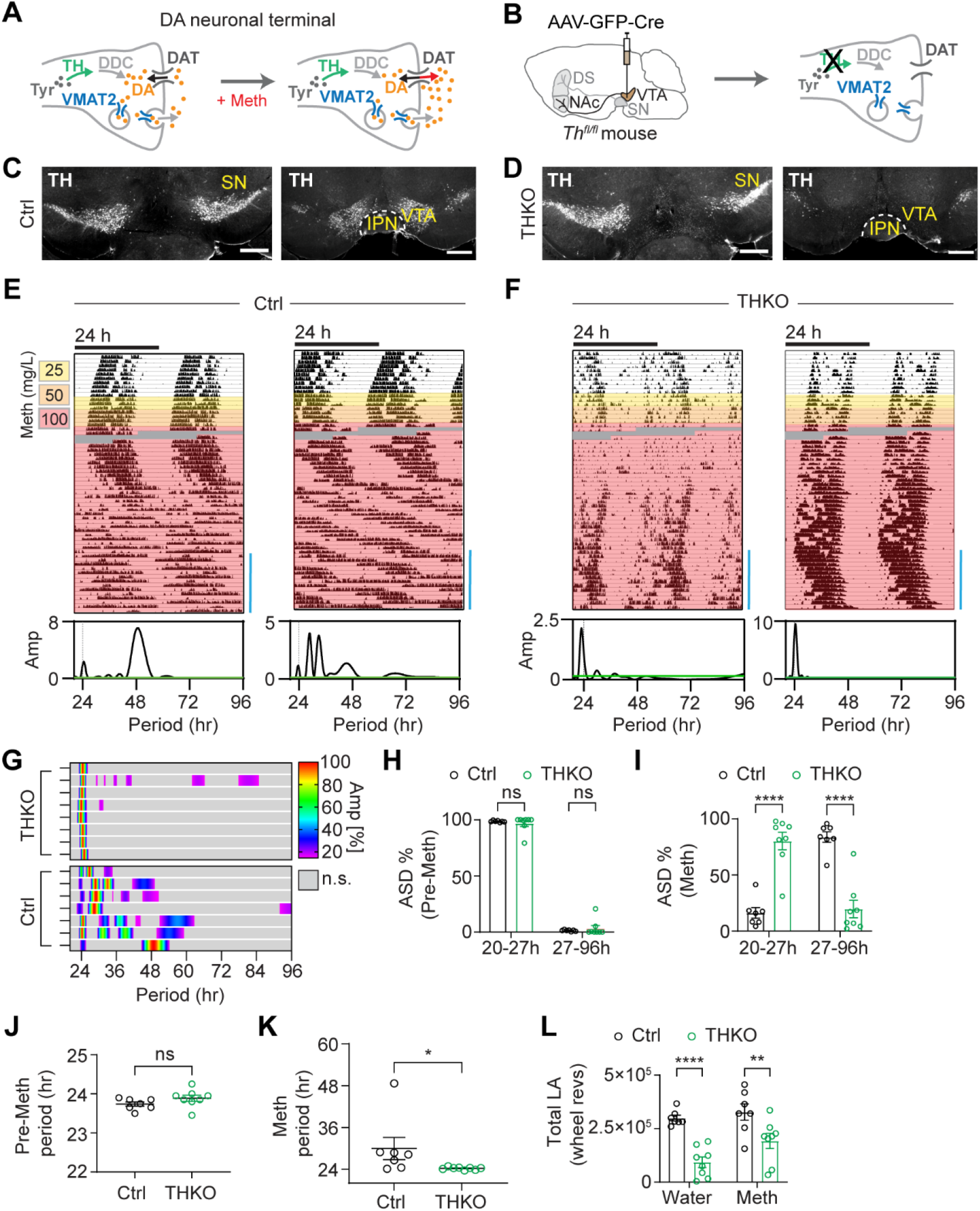
Selective disruption of *Th* in the VTA abrogates infradian rhythm generation capacity. (A) Schematic illustrating the Meth-mediated elevation of extracellular DA at a DA neuronal terminal. DDC, L-dopa decarboxylase; Tyr, tyrosine. (B) Strategy for selective elimination of DA production in DA neurons of the VTA. (C, D) TH immuno-labeling of midbrain sections from saline (Ctrl, C) and AAV-GFP-Cre injected animals (THKO, D). Scale bars, 500 μm. (E, F) Representative actograms showing running wheel activity of control and ^VTA^THKO mice in constant darkness in response to escalating levels of Meth in drinking water. Greyed areas indicate missing data. LS-periodograms correspond to time spans indicated by blue bar. (G) Composite display of normalized periodograms from individual animals corresponding to the final 2 weeks of Meth treatment. (H, I) Periodogram-derived ASD% of significant periodicities in the circadian and infradian period ranges before (H, Pre-Meth) and at the end of Meth-treatment (I, Meth). (J, K) Periodogram-derived highest peak before (J, Pre-Meth) and at the end of the Meth-treatment regimen (K, Meth). (L) Total locomotor activity prior to (Water) and during Meth treatment (Meth). Mean ± SEM. n= 7-8. Mann-Whitney’s test, J and K. Two-way ANOVA with Bonferroni’s multiple comparison, H, I, and L. ns, not significant; \**P*<0.05; \*\**P*<0.01; \*\*\*\**P*<0.0001. See also Figure S3.

### Low locomotor activity does not prevent infradian rhythm generation

Consistent with ^VTA^Casp3, ^VTA^THKO also showed a reduction in overall locomotor activity when compared to controls (Figure 4L), which could be viewed as a contributing factor to the failed 2ndC emergence in response to Meth. To address this possible confounder, we created mice deficient of the vesicular dopamine transporter 2 (Vmat2) specifically in ^VTA^DA neurons, which consequentially abrogates DA vesicular uptake and thus release (Figure 5A).

**Figure 5.**
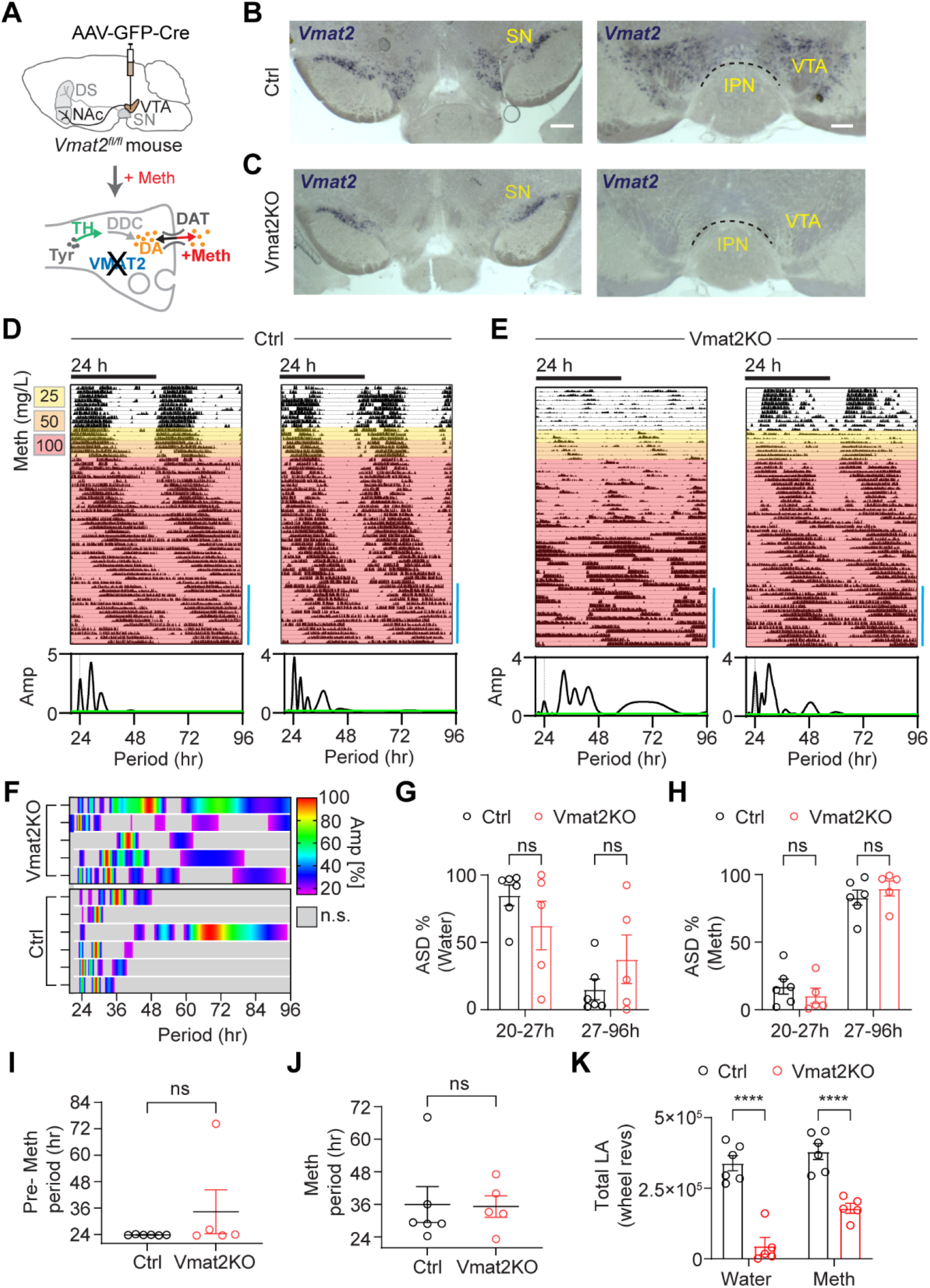
Selective disruption of *Vmat2* in the VTA does not abrogate the capacity for infradian rhythm generation. (A) Strategy of *Vmat2* disruption leading to loss of DA vesicular uptake and release selectively in DA neurons of the VTA. (B, C) Bilateral injection of AAV5-GFP-Cre into the VTA of *Vmat2^fl/fl^*mice leads to loss of *Vmat2* message in the VTA but not the SN. Shown are representative images of brain sections after situ hybridization with a *Vmat2* specific riboprobe of viral (C, VmatKO) and saline-injected mice (B, Ctrl). Blue precipitate indicates *Vmat2* message. Scale bars, 500 μm (SN), 250 μm (VTA). (D, E) Representative actograms displaying running wheel activity of control (D) and ^VTA^Vmat2KO mice (E) in constant darkness in response to Meth in drinking water. LS-periodograms shown below correspond to the time span indicated by blue bar. (F) Composite display of normalized periodograms computed from the final 2 weeks of recording under Meth treatment. (G, H) Periodogram-derived PSD% of significant periodicities in the circadian and infradian period ranges before (G) and at the end of Meth-treatment (H). (I, J) Periodogram-derived highest peak in the 20-96 hr range before (I, Pre-Meth) and at the end of Meth-treatment (J, Meth). (K) Total locomotor activity prior to (Water) and during Meth treatment (Meth). Mean ± SEM. n= 5-6. Mann-Whitney’s test, I and J. Two-way ANOVA with Bonferroni’s multiple comparison G, H, and K. ns, not significant; \*\*\*\**P*<0.0001. See also Figures S4 and S5.

Bilateral injections of AAV-GFP-Cre into the VTA of *Vmat2^fl/fl^* mice led to a preferential loss of *Vmat2* transcripts in the VTA (Figures 5B and 5C), while expression in the SN, raphe nucleus and locus coeruleus remained intact (Figures 5C and S4A, S4B). As was the case for ^VTA^Casp3 and ^VTA^THKO mice, ^VTA^Vmat2KO mice also exhibited a marked reduction in spontaneous locomotor activity (Figure 5K), while the circadian period pre-Meth remained again unaffected (Figure 5I). However, contrary to the other models, ^VTA^Vmat2KO mice retained the ability to produce 2ndCs (Figures 5D and 5E). Both the highest peak (Figures 5I and 5J) and spectral energy distribution (Figures 5F-5H) showed a shift towards infradian periods for both controls and ^VTA^Vmat2KO mice. Similar results were obtained with *Vmat2^fl/fl^*mice that were crossed to *DAT-iCreER* mice ^29^. Tamoxifen-treatment of *DAT-iCreER* x *Vmat2^fl/fl^* mice led to a selective loss of *Vmat2* in DA neurons across the midbrain and a profound loss of locomotor activity (Figures S4C-S4F). However, subsequent exposure to Meth not only increased activity in form of distinct locomotor bouts, but also led to the emergence of 2ndCs (Figure S4F). For further confirmation of the Vmat2KO findings, we injected the VTA of *DAT-Cre* mice with an AAV that expresses tetanus toxin light chain (TetTox) ^30^ Cre-dependently (AAV-DIO-GFP-2A-TetTox) to synaptically silence DA neurons ^30^. TH-positive fibers of the NAc shell were found to comprehensively co-stain for the viral GFP reporter indicating successful targeting of NAc-projecting ^VTA^DA neurons by the virus (Figures S5A and S5B). As with ^VTA^Vmat2KO mice, we found ^VTA^TetTox mice to exhibit reduced locomotor activity at baseline (Figures S5C and S5D), while their 2ndC induction capacity was fully retained (Figures S5C-S5E).

Together, these findings demonstrate that a general, sustained reduction in spontaneous locomotion does not necessarily preclude generation of a 2ndC in response to Meth.

### Antiphasic calcium rhythms in ^NAc^DA-projections

Optogenetic interrogation of DA fibers in major target areas of ^VTA^DA neurons revealed that only stimulation of DA fibers in the NAc promoted wakefulness in mice when sleep pressure is high (early portion of daytime) ^23^, indicating a key role of NAc-projecting DA neurons of the VTA in arousal/wakefulness regulation. We thus wondered if NAc DA fibers in particular would show evidence for rhythms in neuronal activity in line with their role as generators of a second rhythmic rest-arousal component. To isolate the NAc-DA neuronal processes from circadian influences, we crossed *Bmal1^-/-^* mice (Figure 6A) to mice that carried alleles encoding a Cre-activatable fluorescent calcium sensor (GCaMP6s) ^31^ and the *DAT-Cre* transgene to produce clock-deficient mice that express the calcium indicator selectively in DA neurons. These *DAT-Cre x GCaMP6 x Bmal1^-/-^* mice (DAT-G6-BKO mice) were implanted with an optic fiber reaching into the medial shell of the NAc or dorsal striatum to trace fluctuations in cytosolic calcium -as proxy of neuronal activity-exclusively in DA neuronal terminals (Figures 6B and 6C). We then recorded fluorescence changes photometrically alongside locomotor activity using PIR sensors in constant conditions (constant darkness, Figure 6B).

**Figure 6.**
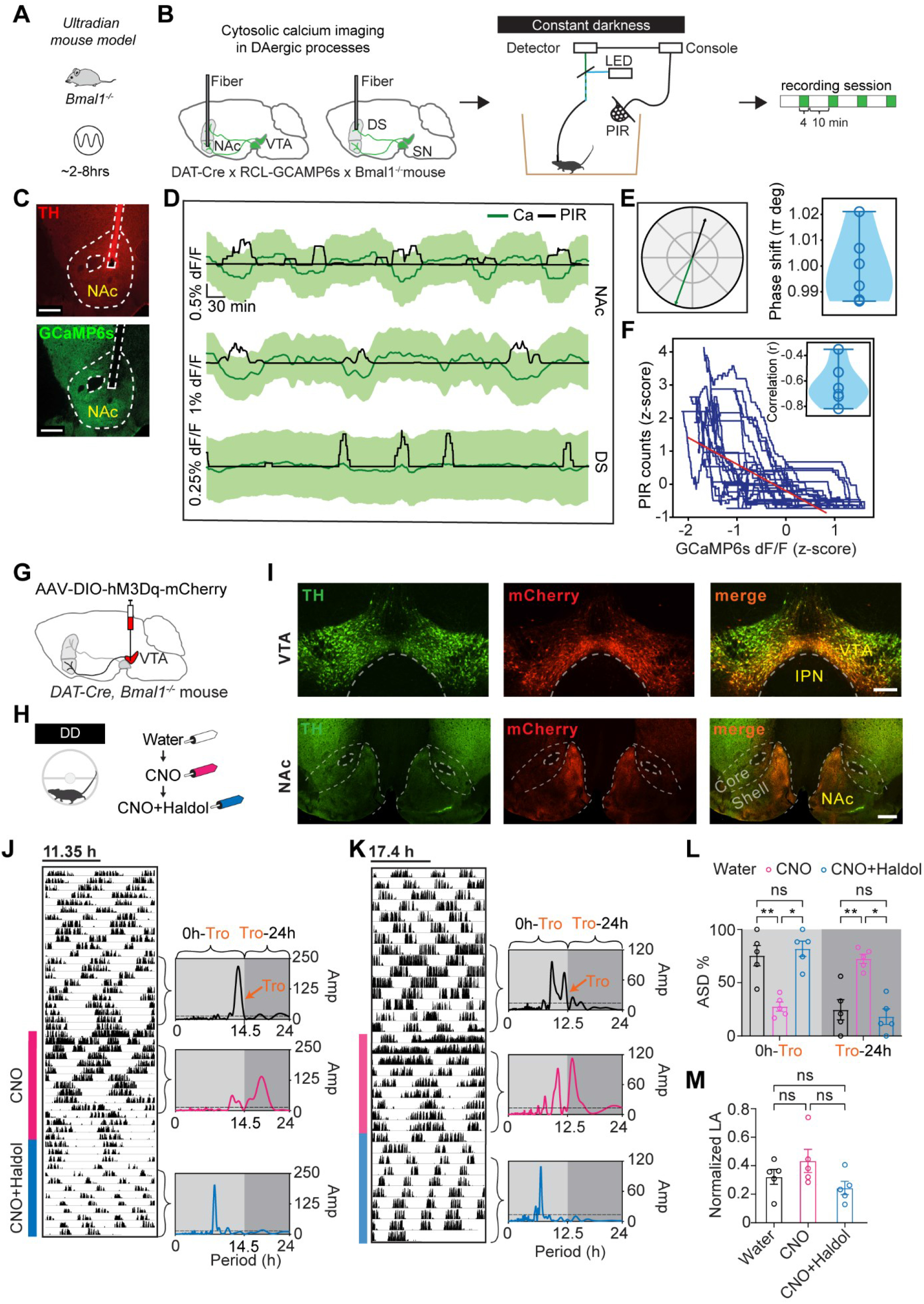
Antiphasic calcium rhythms in ^NAc^DA processes and period shortening of ^VTA->NAc^DA-neuron-driven locomotor rhythms by antipsychotics. (A) Circadian clock-deficient animal model producing ultradian (2-8hr) locomotor rhythms. (B) Experimental layout showing optic fiber targeting of calcium indicator expressing DA neurons and long-term recording of indicator fluorescence at 10 min intervals in constant darkness alongside locomotor activity by PIR. (C) TH and GCamp6s immunofluorescence in the NAc of a reporter mouse with path of recording fiber implant demarcated by dashed line. (D) Example traces of calcium indicator (GCaMP6s) fluorescence (green) and PIR counts (black) from NAc (top and middle) and dorsal striatum (bottom). Shown are smoothened data (10-min average moving window) and standard deviation of the GCaMP6s trace (light green). (E) left: The acrophases of the cosinor fits for the traces shown in c (top recording) are antiphasic; right: Group analysis confirming that the GCaMP6s and PIR signal fluctuations are antiphasic, i.e., shifted by Pi (n= 5, *P* > 0.87 for statistical difference from Pi, one sample t-test). (F) Correlation of GCaMP6s versus PIR traces from recording in c, top. Insert in e: r-values of animal cohort (n = 5, *P* < 0.0001 per animal, Wald Test). (G, H) Experimental regimen of locomotor rhythm monitoring in constant darkness and CNO treatment upon viral delivery of a chemogenetic actuator into DA neurons of the VTA. (I) Immunofluorescence images of VTA and NAc sections labeled for mCherry (red) and TH (green) from *DAT-Cre x Bmal1^-/-^* mice injected with AAV-DIO-hM3Dq-mCherry in the VTA. Core and Shell indicate respective subregions of the NAc. Scale bars, 500 (NAc) and 250 μm (VTA). (J, K) Representative, modulo-plotted actograms showing locomotor responses to CNO in drinking water followed by CNO+Haldol. Periodograms are computed from indicated time windows. ‘Tro’ indicates the *trough* after the dominant periodogram peak during water treatment. The 0hr-Tro periodogram segment therefore captures the periodicities that dominate at baseline (water only treatment). Dashed line, significant threshold for rhythmicity. (L) % of total ASD allocated to 0hr-Tro and Tro-24hr, respectively. (M) Cumulative locomotor activity over 96 hr normalized to the sum of the three 96 hr time windows indicated by brackets. Mean ± SEM. n= 5. Two-way ANOVA with Bonferroni’s multiple comparison. ns, not significant; *P<0.05; **P<0.01.

We found calcium sensor fluorescence from the NAc of DAT-G6-BKO mice to fluctuate in synchrony with their locomotor activity rhythm seemingly adopting an antiphasic relationship (Figures 6D-6F). By contrast, such pronounced fluctuation in reporter fluorescence was not observed when recorded from the dorsal striatum (Figure 6D, bottom trace). Cosinor analysis of the NAc-recordings validated fluorescence signal rhythmicity (n=7, fit significance was q<0.001 across animals). In line with the antiphasic relationship, locomotor activity versus calcium-reporter phase offset was consistently indifferent from pi (one-sample t-test, *P*>0.87, Figure 6E) and locomotor bouts versus GCaMP6s signal was also found to be negatively correlated by regression analysis (Figure 6F). These results indicate an intrinsic capacity of ^NAc^DA-processes to produce oscillations that are synchronized to locomotion, which supports a role of these processes also in infradian rhythm generation.

### Period-lengthening by chemogenetic activation of NAc-projecting DA neurons is counteracted by antipsychotic treatment

To assess if specific activation of NAc-projecting ^VTA^DA neurons is sufficient for period lengthening, we injected *DAT-Cre x Bmal1^-/-^* mice with the Cre-activatable AAV-DIO-hM3Dq-mCherry^32^ into the ventral VTA to drive expression of the chemogenetic activator hM3Dq specifically in TH neurons of the VTA (Figures 6G and 6H). The mCherry expression pattern confirmed the expected preferential transduction of TH neurons in the ventral VTA and corresponding processes in the NAc medial shell (Figure 6I). Addition of the hM3Dq-ligand CNO to the drinking water lengthened locomotor period (Figures 6J-6L). Next we applied haloperidol (Haldol), a common antipsychotic and dopamine receptor D2 inverse agonist ^33^, which we have previously shown to counteract Meth-mediated period lengthening in clock deficient mice but also 2ndC emergence in clock intact mice ^12^. Haldol supplementation reversed the CNO-mediated effect on locomotor period in the hM3Dq expressing mice (Figures 6J-6L) suggesting that CNO acts on the same oscillatory process as Meth treatment or DAT disruption does. Notably, neither CNO nor CNO+Haldol treatment significantly altered total locomotor activity (Figure 6M), suggesting that the change in period is not a mere consequence of a change in overall activity.

### Infradian rhythm emergence requires NAc-projecting DA neurons

The chemogenetic and fiberphotometry findings support a role of NAc-projecting DA neurons in non-circadian rhythm generation. We therefore wondered if these neurons are *necessary* for infradian rhythm emergence and thus injected the catecholaminergic neurotoxin 6-hydroxy-dopamine (6-OHDA) bilaterally into the NAc medial shell region to selectively ablate DA neurons that project to the NAc (Figure 7A). Immunohistochemical examination of ^NAc^6OHDA mice showed the expected preferential loss of TH signal in the NAc and corresponding reduction but not complete elimination of the TH immuno-signal in the VTA (Figures 7B and 7C), as expected given that the NAc is only one of several ^VTA^DA neuronal targets^34^. Mirroring the behavior of ^VTA^Casp3 and ^VTA^THKO mice, ^NAc^6OHDA animals largely failed to produce a 2ndC in response to Meth (Figures 7D-7H, and S6A, S6B). In line with the other models, the circadian period pre-Meth remained unaffected (Figure 7G), while locomotor activity tended to be lower under pre-Meth conditions (Figure S6C).

**Figure 7.**
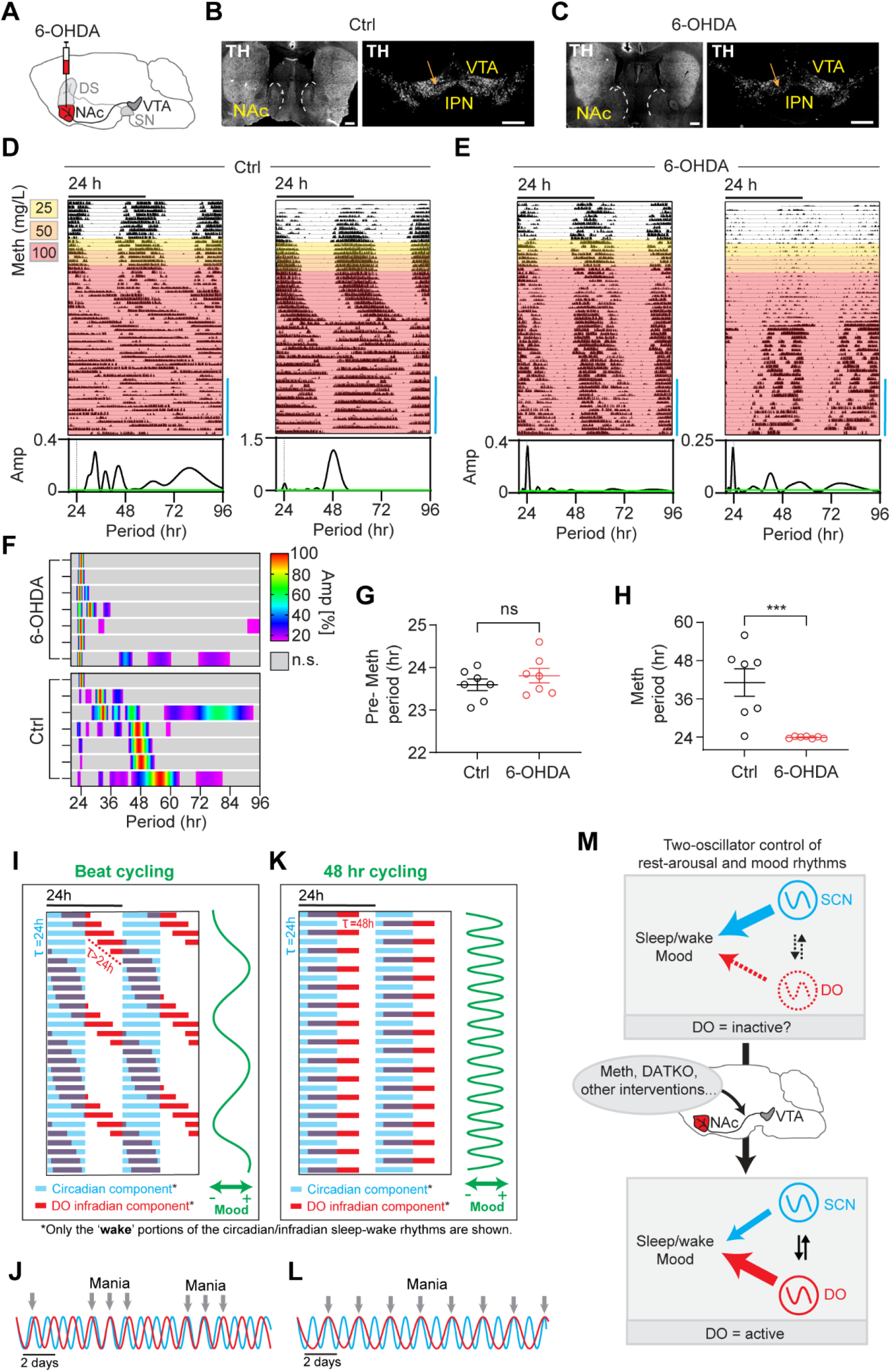
Loss of infradian rhythm generation capacity upon NAc-projecting DA neuronal ablation and two-oscillator model for BD cycling. (A) 6-hydroxydopamine (6-OHDA) was bilaterally injected into the NAc to ablate DAergic processes. (B, C) TH immunolabeling of striatal and midbrain sections from saline (B) and 6-OHDA (C) injected animals. Scale bars, 500 μm. (D, E) Representative actograms showing running wheel activity of saline (D) and 6-OHDA-injected mice (E) in constant darkness in response to escalating doses of Meth in drinking water. LS-periodograms correspond to the time interval indicated (blue bar). (F) Composite display of normalized periodograms from individual animals computed from the final 2 weeks of recording under Meth treatment. (G, H) Periodogram-derived highest peak in the 20-96 hr range before (G, Pre-Meth) and at the end of Meth-treatment (H, Meth). (I) Model actogram showing daily wake periods reflective of a DO operating at a frequency non-harmonious to the SCN clock. (J) In-and-out of phase ‘beating’ of the DO and SCN clock when frequencies are non-harmonious resulting in mood cycling. (K, L) Model of sleep-wake and mood cycling at a DO period of 48 hr which is in harmony with the 24 hr SCN clock. (M) Upon ‘activation’, the DO influences/dominates sleep-wake and mood rhythms. Mean ± SEM. n= 7. Mann-Whitney’s test. ns, not significant; ***P<0.001. See also Figure S6.

## Discussion

We previously showed that global elimination of DAT can lead to the emergence of a 2ndC ^12^, i.e., a manipulation that selectively yet broadly targets DA neurons is sufficient for infradian rhythms induction. Our new data now demonstrate the necessity of NAc-projecting DA neurons of the VTA for infradian rhythm emergence arguing for a role as infradian rhythm generators.

### Relevance of DA, but not its cellular release mode for 2ndC expression

Both Meth treatment and DAT knock out result in 2ndC emergence and elevation in extracellular DA, yet the latter seems to be achieved by different mechanism: a reverse-acting DAT in case of Meth and vesicular DA release without reuptake in case of *DAT^-/-^*mice. The preservation of Meth-induced infradian rhythmicity in the absence of DA-vesicular release in (*Vmat2KO*) or general synaptic silencing (Tetox experiment) further underscores that while DA release by ^VTA^DA neurons is of critical importance for 2ndC induction, the actual release mode is not. The Tetox-silencing experiment also indicates that neurotransmitter release in general by ^NAc^DA terminals is dispensable for infradian rhythm generation.

### Calcium fluctuations in ^NAc^DA processes as cellular correlate for behavioral rhythms

The antiphasic relationship between cytosolic calcium fluctuations in ^NAc^DA fibers and locomotor rhythm presented here contradicts our previous findings of a phasic relationship between extracellular DA in the striatum/NAc region and locomotor rhythms^12^. Furthermore, an increase in intracellular calcium is generally considered to be an indicator of increased neuronal activity^35^ and thus synaptic transmitter release and accordingly, intracellular calcium should peak in phase with extracellular DA and locomotor activity. However, in our study, the calcium fluctuations observed are on a very different timescale and resolution when compared to previous studies that found correlates between neuronal firing and cytosolic calcium levels (see for instance ^35^). In this context it is interesting to note that a robust phase offset between cytosolic 24 hr calcium rhythms and neuronal firing was previously shown for SCN explants in culture ^36^, further corroborating that slow cytosolic calcium rhythms may not directly read out changes in neuronal electrical activity.

### MASCO vs DUO vs DO

Our finding that the period of 2ndC reaches far into infradian range challenges the currently held view that the oscillator which Meth engages is circadian in nature, a notion also reflected in the wide-spread use of ‘MASCO’ (methamphetamine-sensitive circadian oscillator) as descriptor for this process ^7,13^. Considering our previous findings that 2ndC emergence does not require Meth but instead can also result from DAT disruption^12^ along with our data presented here (Figure 1), we now propose to use ‘*dopaminergic oscillator*’ or DO to identify the oscillator that drives infradian rest-arousal cycles and relies on DA production. This more general descriptor should also take precedence over the use of the term “dopaminergic ultradian oscillator (DUO)” we previously coined for this oscillator^12^, given the wide range of periods it can adopt.

### BD relevance of 48hr rhythmic mice

48 hr rhythms in rest-arousal could be considered a peculiar finding due to the scarcity of its accounts in the animal and human literature^13,37^. However, a recent BD case study showed that pausing treatment of the widely used antipsychotic aripiprazole resulted in the emergence of 48hr cycles in sleep-wake with sleep cycles again normalizing to daily (24 hr) rhythms after drug reinstatement^38^. This demonstrates that standard medication can obscure the propensity of cycling in BD which in turn suggest that 48 hr cycling may be much more widespread in BD than currently acknowledged.

In further support of 48hr rhythmic mice representing a valid model for cycling in BD, we found them to display hyperactivity and more straight path movements on days with short sleep. While these data indicate face validity for cyclical BD mania, there is also evidence for predictive validity as antipsychotic treatment corrects infradian rhythmicity in both Meth and DATKO models much in line with the above mentioned case report ^12^. Finally, high-dose methamphetamine has been recently reported to trigger bipolar mood cycling de novo ^39^, which can be viewed as evidence for construct validity.

These accounts together with the fact that the dopamine system is one of the principal BD treatment targets ^40^ argue that the 48 hr rhythmic mice characterized here represent a bona fide model for 48 hr cycling in BD. This in turn would then suggest that NAc-projecting DA neurons not only serve as substrate for murine infradian rhythm production, but also 48 hr cycling in BD. While there exists a considerable number of mouse models for mania or depression^41-43^, animal models for cycling or mood switching are scarce^44^. Yet understanding state switching mechanistically is considered the ‘holy grail’ of bipolar disorder^45^. 48 hr cycling mice may therefore critically help elucidate the cellular and molecular basis of this disorder.

### Towards a general mechanistic understanding of bipolar cycling

Mood switching in BD with great regularity has been also reported at periodicities much longer than 48 hr, such as weeks or even months ^46-48^, and it has been already proposed that this BD rapid cycling could be due to the periodic in- and out-of-phase ‘beating’ of two rhythmic processes that deviate in their frequencies ^49^. Given what we know about the DO, it seems very plausible that this oscillator and the circadian clock represent those rhythmic processes with the beat frequency and thus mood switch frequency determined by the period of the DO, which is uniquely tunable. For instance, with the circadian timer set at ∼24 hr and the DO at ∼27 hr, the beat frequency would be 13 days, i.e. the DO and circadian timer would be in phase every 13 days. If phase alignment associates with (hypo)mania, then this setting would produce recurrent mania with a 13-day period. Therefore, the period of the DO would determine the inter-episode interval of cycling subjects, while the degree of relative coordination with the circadian clock may determine the length of the manic episode (Figures 7I-L). The sinusoidal cycling of mood in phase with sleep phase cycling as observed in one renown BD case report^48^, further corroborates the proposed beating mechanism.

Our findings specifically support a role of NAc-projecting DA neurons in the cycling aspect of BD, where they act as drivers of mood switches via in- and out-of-phase alignment of the DO with the circadian clock/external light cycle. In this context it is intriguing to note that a recent positron emission tomography (PET) imaging study using a DAT ligand as tracer revealed lower DAT availability in manic subjects versus controls selectively in striatal regions including the NAc^50^. As the DAT is exclusively expressed by DA projections in the NAc, this data provides further credence for the DA projections of the NAc as site or drivers of cyclicity in BD. In line with this recent PET data, diffusion tensor imaging of a rapid cycling female subject who exhibited 2-weeks of mania in association with each menstrual cycle, showed increased fractional anisotropy in the manic state which was largely restricted to the NAc region when compared to the subject’s preceding and succeeding euthymic states, or to unaffected control subjects^51^. This latter study can be viewed as first evidence for a correlate of BD cycling in human brain activity which intriguingly centers on the NAc, in further support of NAc-projecting DA neurons as drivers of infradian rhythms and bipolar cycling.

### DO state in unchallenged mice and healthy humans

While we provide evidence that the DO can produce infradian rhythms at a wide range of periods upon manipulation of the DA system, there is the question about the DO’s state in intact laboratory rodents without a challenge und in healthy humans. While the DO may adopt non-infradian periodicities under such conditions, it could as well be inoperative with temporal sleep-wake regulation solely resting on the circadian timer. In such scenario, the DO would constitute an ‘activatable’ rhythm generator that can jump into action when challenged by e.g. psychostimulants or in bipolar disorder subjects (Figure 7M). Due to its robust rhythm generation capacity, it seems very likely that the DO has functions beyond the pathological context -whether as a constitutive active or activatable entity-, which however have yet to be elucidated.

### Limitations of the study

Locomotor activity-linked rhythms of intracellular calcium in DA-projections of the NAc were detected in circadian clock-deficient mice, which were not exposed to Meth and thus only exhibited ultradian rhythms in rest:arousal. While this finding demonstrates a principal capacity of intracellular rhythm generation in DA processes of the NAc, calcium indicator recording upon chronic Meth exposure would be required to link calcium fluctuations to infradian rhythmicity/48 hr cycling. Additionally, while our study demonstrates the necessity of ^NAc^DA processes for infradian rhythm generation, it is not yet clear if these processes are mere components of the DO or harbor it in its entirety much like the circadian clock is contained in SCN neurons.

## Acknowledgements

We thank M. Jaramillo Garcia and the Molecular and Cellular Microscopy Platform at the DHRC for assistance with fluorescence imaging, S. Srikanta for help with data analysis, F. Tronche for the *DAT-Cre* mice, and the drug supply program of the National Institute on Drug Abuse for providing methamphetamine. This work was supported by grants from the Canadian Institutes for Health Research (PJT-148914 and PJT-487638, K.F.S.; PJT-148790, M.V.K.; PJT-180590, P.V.S.; 201309OG-312343-PT and 201803PJT-399980-PT, B.G.), Natural Sciences and Engineering Research Council of Canada (RGPIN-2020-06610, K.F.S.; RGPIN-2022-03390, P.V.S.), National Institutes of Health (R01-AG062514-03 and R01-AG067193-02, M.D.), and the Graham Boeckh Foundation (B.G.).

## Author contributions

K.-F.S., P.S.M., and C.B. conceived and designed the experiments with contributions by M.V.K and B.G. P.S.M, C.B., and L.Z. performed, and K.-F.S. supervised the experiments. B.G., and M.D. produced and provided resources. All authors contributed to data analysis and interpretation. P.S.M., C.B., and K.-F.S. wrote the manuscript.

## Methods

### Animals

*Bmal1^−/−^* mice ^52^ were crossed to *DAT-Cre* mice ^25^ to enable viral targeting of DA neurons, or additionally crossed to Ai96(RCL-GCaMP6s) ^53^ for fiber photometry recordings. *Th^flox/flox^* mice ^28^ were used for viral-mediated *Th* disruption in the VTA. *Slc18a2^flox/flox^* mice ^29^ were virally targeted for midbrain *Vmat2* disruption or crossed to *DAT-iCreER* (JAX stock #016583) mice for body-wide, DA neuron-specific *Vmat2* disruption during adulthood. Mouse lines were on a C57BL/6 background. All animal procedures were carried out in accordance with the recommendations of the Canadian Council on Animal Care (CCAC) and have been approved by the local McGill University Animal Care Committee.

### Locomotor activity monitoring and sleep analysis

Animals were individually housed in running-wheel cages in light-tight cabinets in constant darkness. Locomotor activity was recorded continuously using Clocklab data collection software (ClockLab, Actimetrics, Wilmette, IL). ClockLab analysis software was used to generate actograms displaying binned running wheel revolutions per 5 min (0.12 hr), Lomb-Scargle ^54^ periodograms as well as heatmap displays of Morlet Continuous Wavelet Transforms and associated wavelet ridge plots.

Passive infra-red: Animals were individually housed in standard cages. Locomotion was recorded at 0.25Hz and binned at 1-min intervals using custom software. Locomotion data was transformed into actogram displays using MATLAB (The MathWorks) custom scripts.

Sleep counts were generated by assigning a score of 1 to any 1-min bin with a value of 0, in concordance with ^17,18^. For sleep analysis during 48 hr cycling, we selected 4-day time spans of 48 hr rhythmicity (Figure S1) for each animal using spectral power quantification of circadian (22-26 hr) versus infradian (44-52 hr) range frequencies as guide for selection.

### In situ hybridization

In situ hybridization was performed as described previously ^55^. Briefly, fixed brains collected after intra-cardiac perfusion (Z-fix: 10% aqueous buffered zinc formalin, Anatech LTD, Battle Creek, Michigan, USA) were cut at 25 microns using a cryostat (Leica, Solms, Germany) and stored at −80°C until hybridization. Sections were hybridized overnight at 60°C to a digoxigenin-labeled riboprobe targeting the coding regions of mouse *Vmat2 (*nucleotides 141– 255 of the *Slc18a2* mRNA, Genbank, NM_172523.3).

### Immunohistochemistry

Immunostaining was performed as previously described ^56^. Briefly, mice were deeply anesthetized and transcardially perfused with 10% formalin (Z-fix, Anatech). Brains were postfixed in 10% formalin overnight and then incubated in 30% sucrose in saline for at least 24 hours. Brains were cut at 40 microns using either a cryostat (Leica, Solms, Germany) or vibratome (VT 1200S, Leica, Wetzlar, Germany) and collected in 3 or 4 series per brain, respectively. For immunohistochemistry, sections were rinsed in PBS (pH 7.4), incubated in blocking solution (1% goat or donkey serum in 0.3% Triton X-100 in PBS) for 1 hr, followed by incubation with the primary antibody in blocking solution at 4^°^C overnight. Sections were then incubated in secondary antibodies for 2 hours at RT. Sections were mounted on superfrost slides (VWR, Radnor, PA), coverslipped with Vectashield mounting medium with DAPI (Vector Labs, Burlington, Canada) and imaged by fluorescence microscopy. Primary and secondary antibodies were used at the following dilutions: rabbit anti-RFP, 1:1000 (Rockland, Limerick, PA) to enhance detection of mCherry expression, anti-GFP, 1:1000 (Invitrogen, Life Technologies, Carlsbad, CA) for GCaMP6s detection and mouse anti-TH, 1:1000 (EMD Millipore, Etibicoke, Canada) for tyrosine hydroxylase detection; Alexa 488 and Alexa 568 conjugates (Life Technologies) were used as secondary antibodies at a dilution of 1:500.

### Pharmacology

Tamoxifen: To activate Cre recombinase, *DAT-iCreER x Vmat2^fl/fl^*mice were i.p. injected with tamoxifen (Sigma; dissolved in a 1:9 (v/v) ethanol:sunflower oil mix to a final concentration of 30mg/mL) twice daily for 5 consecutive days. The first dose (0.6mg) was given 3 hrs after lights on and the second (0.75mg) 1 hr before lights off.

Methamphetamine, Haloperidol, Clozapine-N-oxide (CNO): Stock solutions of (+)-Methamphetamine hydrochloride (25-100 mg/l, National Institute of Mental Health, Bethesda, MD), Haloperidol (11.25 mg/ml, Sigma-Aldrich) and Clozapine-N-Oxide (15 mg/l, National Institute of Mental Health, Bethesda, MD) were prepared using drinking tap water. Haloperidol was dissolved by stirring at 40°C. Drugs were added to the drinking water to reach concentrations as indicated. For i.p. injection, a methamphetamine hydrochloride solution was prepared using saline and injected at a dose of 2.5 mg/kg.

### ASD and CWT computations

To determine the prevalence of oscillations in a given period range we calculated the sum of amplitude of all significantly rhythmic periodicities (α = 0.001) in the Lomb-Scargle periodogram, that is, the power spectral density. Densities for circadian (20-27 hr) or infradian (27-96 hr) ranges were then normalized by division with the total significant spectral density (20–96 hr) and expressed as percentage according to the following formula:

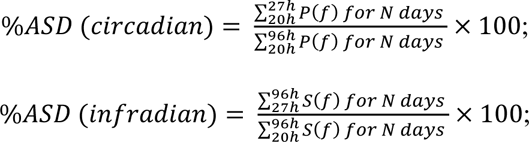

where *P*(f) represents the power calculated by the Lomb-Scargle method ^54^.

Continuous wavelet transforms and scale-averaged wavelet spectra were computed using the MATLAB Wavelet Toolbox with Morse wavelets and 24 voices per octave. The windows used for computing the scale averaged wavelet spectrum were 20-27 hr and 27-72 hr, respectively. To quantify the effect of chemogenetic activation in AAV-DIO-hM3Dq-mCherry^32^ injected *DAT-Cre x Bmal1^-/-^* mice on locomotor period, changes in power distribution were calculated as follows: first we identified the dominant peak in the periodogram computed from the 96hrs timespan prior to CNO treatment. We then used the trough that followed the dominant peak as reference point for quantifying shifts in power distribution (indicated by Tro in Figure S6). Accordingly, we calculated the % of total ASD allocated to the periodogram segments ‘0 hrs to Tro’ and ‘Tro to 24hrs’ of this trough, respectively. The ‘Tro’ can be considered analogous to the 27 hr mark dividing the periodogram in circadian (20-27 hr) versus infradian (27-96 hr) ranges in the context of Meth exposure in circadian intact mice. Note that -in contrast to the circadian period in intact animals-the ultradian locomotor period profoundly varies among *Bmal1^-/-^* animals, typically in the 2-8hr range. Therefore, a fixed reference point such as 27hr in case of circadian intact mice cannot be used and hence the use of Tro whose position is dictated by the ultradian period/spectral power, which is specific to each *Bmal1^-/-^*mouse.

### Virus and 6-OHDA injections

Mice were anaesthetized with isoflurane and placed in a stereotaxic apparatus (David Kopf Instruments). Recombinant AAV vectors were bilaterally injected into to the VTA area (coordinates from bregma: AP: −3.44 mm, DV: −4.40 mm, L: ±0.48 mm; through a cannula (33 gauge, Plastics One) at a flow rate of 0.1 μl/min for 3 min (0.25-0.3 μl total volume per side) using a syringe pump (Harvard Apparatus). Mice were subsequently maintained in individual housing for at least 2 weeks prior to CNO treatment and/or locomotor activity recording. Viruses: AAV5-hSyn-GFP-Cre (University of North Carolina (UNC) Vector Core, titer 3.5x10^12^ genomes copies per ml, diluted 1:1 with saline); AAV5-Flex-taCasp3-TEVp (9) (UNC Vector Core, titer 4.6x10^12^ gc/ml); AAV8-hSyn-DIO-hM3D(Gq)-mCherry (14) (MTP, CERVO Brain Research Center, Laval, titer 7.9x10^12^ gc/ml); AAV8-hSYN1-DIO-GFP-2A-TetTox (University of Michigan Vector Core, titer 8.22x10^13^ gc/ml). The AAV-hSYN1-DIO-GFP-2A-TetTox plasmid was generated by PCR amplifying the GFP-2A-TetTox cassette (from addgene plasmid #166603, gifted by Martin Myers Jr) and introducing AscI and BsiWI enzyme sites to the 5’ and 3’ ends, respectively. Following enzyme digestion, the GFP-2A-TetTox insert was ligated into a AAV-hSYN1-DIO backbone (from addgene plasmid # 44361, gifted by Bryan Roth) which had been digested with AscI and BsrGI.

6-OHDA (10ug/ul concentration) was injected in the medial NAc region (coordinates from bregma: AP: +1.10 mm, DV: −4.40 mm, L: ±0.50 mm), at a flow rate of 0.1 μl/min for 2.5 min (2.5ug per side), equal volume of saline was injected in the control group. Each animal received 25mg/kg Desipramine hydrochloride i.p., 30 mins prior to the surgery to spare norepinephrine neurons from 6-OHDA-mediated neurotoxicity.

### Fluorescence fiber photometry

A 5 mm long optic fiber with a 40 μm core and a numerical aperture of 0.57 was implanted into the NAc (AP: +1.2mm, ML: 0.45mm, DV: -4.6mm) of animals expressing GCaMP6s in DAT positive neurons to deliver exciting light to GCaMP6s and collect fluorescence. The animals were allowed to recovery from implantation surgery over 3 weeks. Changes in GCaMP6s fluorescence in DA projections were assessed using a 1-site 2-color fiber photometry system from Doric Lenses. Calcium sensitive fluorescence was assessed using a 490 nm light source oscillating in power intensity from 0.4 to 1V at 208.616 Hz and isosbestic fluorescence was assessed using a 405 nm LED light source oscillating at 572.205 Hz between 0.4 and 0.8V. Emitted light was measured with a 2151 Femtowatt Photoreceiver and digitized at 12kHz and this signal was software-demodulated according to the two carrier frequencies and low-pass filtered at 12 Hz. Fluorescence was recorded for up to 10 hours for continuous recordings, or for 4min every 6 to 11min for longer recordings. To account for movement artifacts, the isosbestic control signal was then scaled using a least-square linear regression to best fit the 490 nm calcium-sensitive signal and then subtracted from the 490 nm signal. The dF/F percentage was obtained by dividing these corrected values by the 405 nm signal and multiplying by 100. For recordings exhibiting a strong initial nonlinear decay, the first hour was removed from the data. If necessary, a polynomial of degree 1 to 3 depending on the length of the recording was then fitted to the dF/F so the overall shape of the signal was flat. Finally, a 10-min average moving window was applied.

### Open field test and video analysis

The open field test (OFT) was carried out in a transparent box (50cmx50cm floor space) over 10 minutes per animal at ZT0-6 or ZT12-18 in mice with no Meth and at CT0-6 in Meth exposed mice exhibiting 48hr cycling. All experiments were carried out under constant dim light conditions and Camera Module HBVCAM-F20216HD (Walfront Store) was used to video record the behavior at 30fps. The location tracking module of ezTrack ^57,58^ was employed to extract the animal position (x-y coordinates) and total distance travelled.

### Spatial d computation

Spatial d was calculated based on fractal analysis as described before^59^. Briefly, the animal’s spatial trace (derived from the frame-by-frame spatial displacement) extracted from the 10mins OFT recording was divided into 2 cm segments and only the first frame of each of the 2 cm segment associated video frames were retained. We considered these frames as ‘microevents’ in accordance to Paulus and Geyer (1991) and used them for further analysis. We then measured the animal’s spatial displacement at different resolutions k (1, 2, 4 and 8), to compute the total length (L_k_) of the trace at each resolution. Thus, L_k_ is the sum of the distances between the 1^st^and k+1^th^microevent followed by distance between k+1^th^ and 2k+1^th^ microevent and so forth. Next, log_2_(L_k_) vs log_2_(k) was plotted and a non-linear curve fit using least-square regression was applied (GraphPad), with spatial d representing the (-) slope of this fit.

### Statistical analysis

Data are presented as mean ± standard error of the mean. One-way and repeated measures, two-way ANOVA followed by a Bonferroni post hoc test, Student’s t-test, Mann-Whitney U test, Wilcoxon’s test, Wald test, F test, and linear regression analyses were used as indicated in figure legends. Cosinor analysis was performed using the function test_cosinor_pairs from the Python package CosinorPy ^60^.

### Data availability

The data that support the findings of this study are available from the corresponding author upon reasonable request.

### Code availability

Custom computer codes used in this study will be made available upon reasonable request.

## Supplemental figure titles and legends

**Figure S1.**
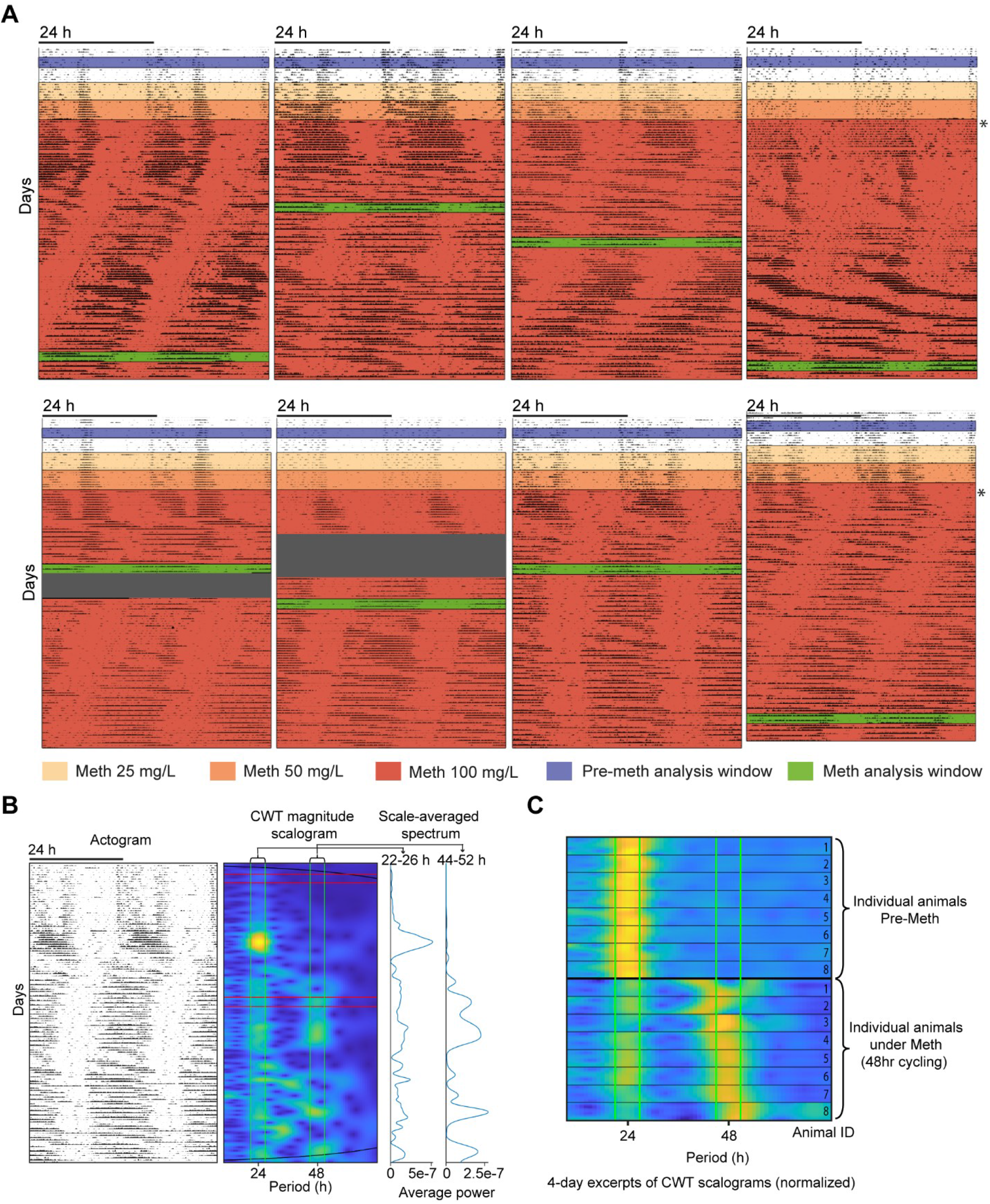
Meth-induced 48-hour rhythmicity in general cage activity. Related to Figure 2. (A) Representative actograms displaying locomotor activity of single-housed animals recorded by PIR sensors. Note that every animal of the cohort tested (N=8) was able to produce a 48hr rhythmic component in response to Meth supplementation of the drinking water. Asterisk (right next to actograms) marks transition into constant darkness. (B) Selection of time spans for sleep analysis during 48hr cycling. Representative actogram showing locomotor activity from PIR data, along-side the CWT scalogram and scale-averaged spectral power for the circadian (22-26hr) and infradian (44-52hr) period ranges (also demarcated by green lines in the scalogram). Red lines demarcate the 4d-time span used at baseline (pre-Meth) and during Meth treatment. Selection of the second time span (Meth) was guided by the presence of a meaningfully strong peak in the 44-52hr spectral power trace. (C) Group display of selected time spans used for analysis.

**Figure S2.**
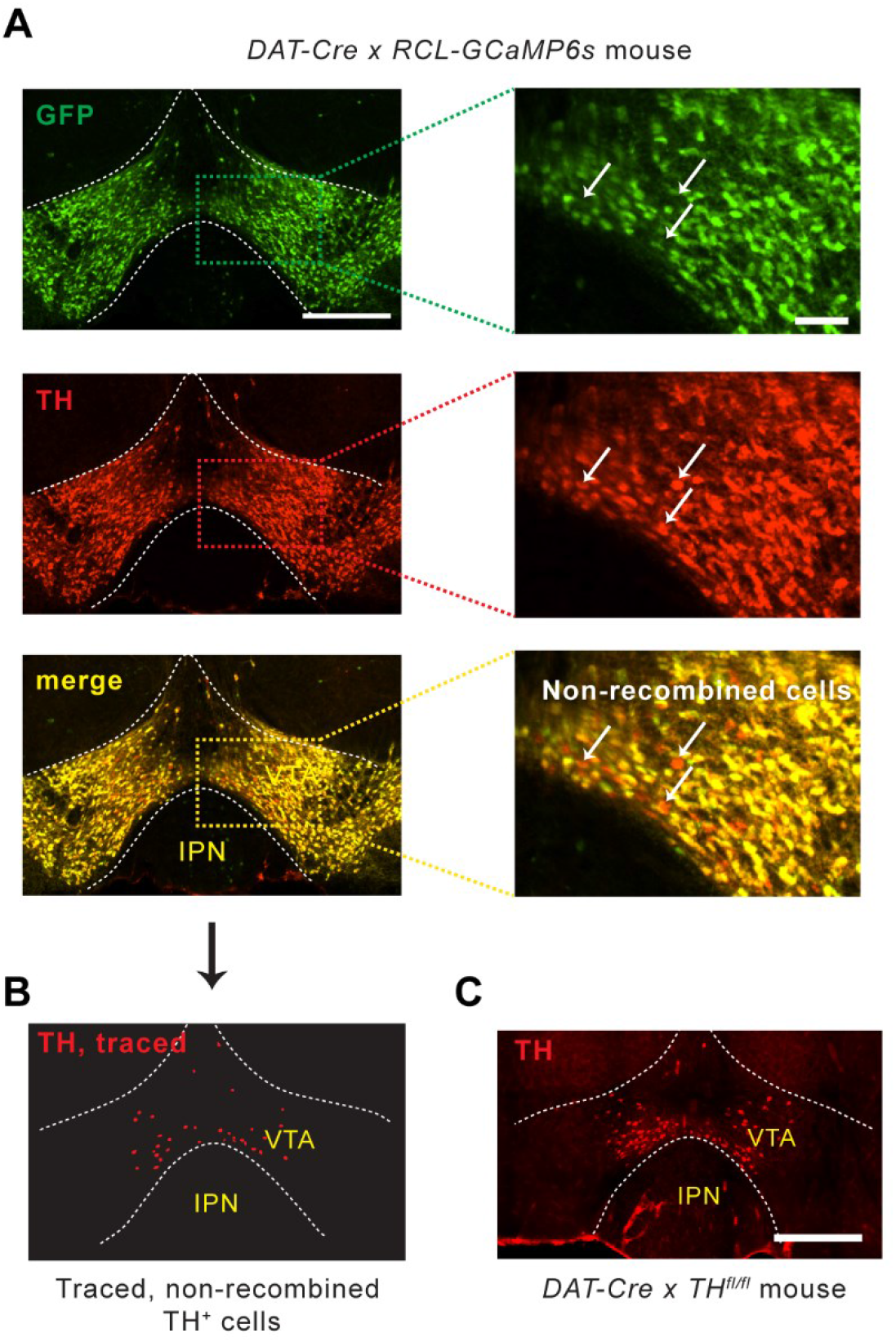
Assessment of Cre activity in the VTA of the *DAT-Cre* mouse line. Related to Figure 3. (A) Some of the TH+ cells that are preferentially located in the ventral VTA apical to the IPN were found to lack GCaMP6s expression in *DAT-Cre x RCL-GCaMP6s* mice. Shown are immunofluorescence images of the VTA stained with antibodies against TH and GFP, respectively. Images on the right show enlargements of the boxed areas of the images on the left. (B) Selective display of non-recombined (GFP-negative) cells that were manually identified in (A). (C) TH immunostaining in the VTA of a *DAT-Cre x TH^fl/fl^* mouse. IPN, interpeduncular nucleus. Scale bar, Scale bars, 500 μm (A, left and C) and 100 μm (A, right).

**Figure S3.**
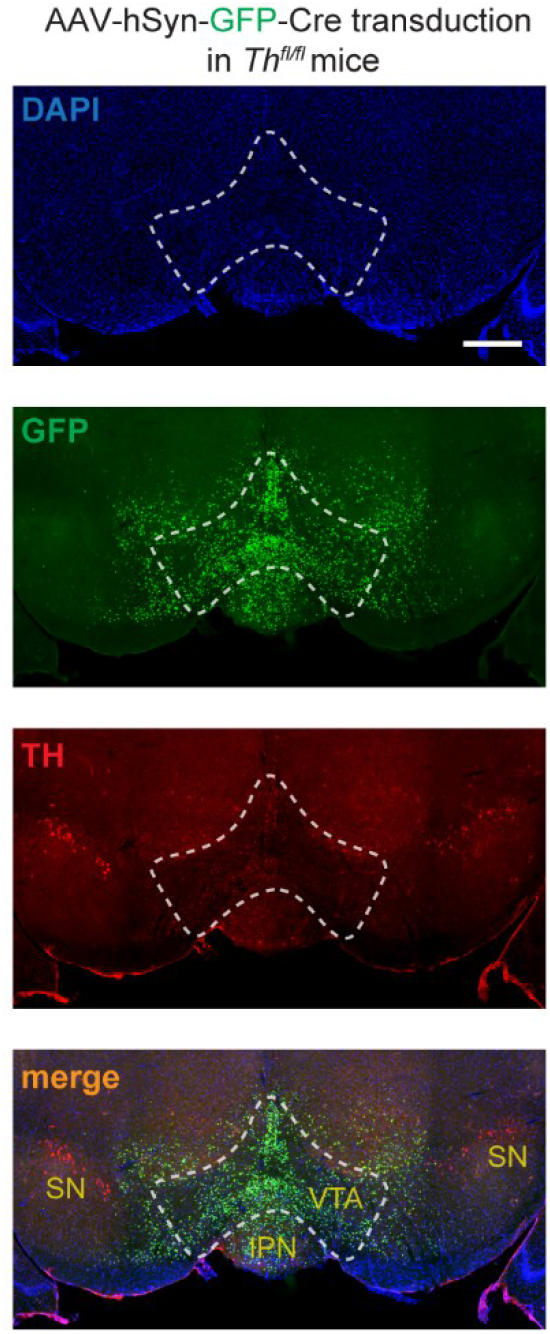
AAV-GFP-Cre viral spread is limited to the VTA region. Related to Figure 4. Midbrain GFP/TH immunofluorescence after bilateral injection of AAV-GFP-Cre into the VTA of a *Th^fl/fl^* mouse. The nuclear GFP signal spread indicates that viral transduction was largely limited to VTA, sparing the SN. Scale bar, 500 μm.

**Figure S4.**
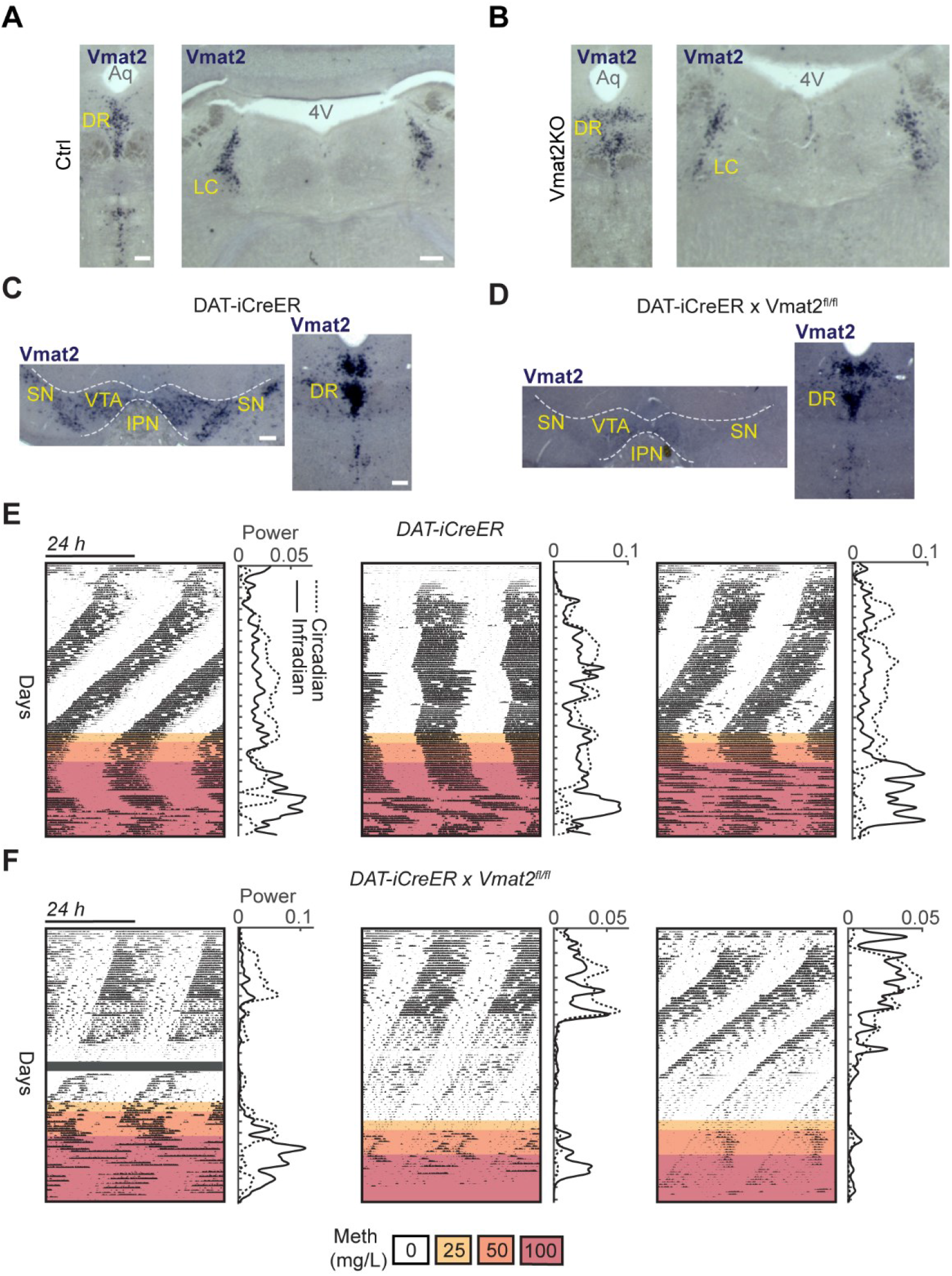
*Vmat2* gene disruption and infradian rhythm generation. Related to Figure 5. (A, B) In situ hybridization with a *Vmat2* riboprobe showing preservation of *Vmat2* signal in raphe nucleus (left) and locus coeruleus (right) in ^VTA^Vmat2KO mice (B) versus controls (A). Scale bars, 500 μm (LC), and 100 μm (Raphe). DR dorsal raphe; LC, locus coeruleus. (C, D) In situ *Vmat2* expression analysis of *DAT-iCreER* (C) and *DAT-iCreER x Vmat2^fl/fl^* (D) mice upon tamoxifen treatment revealing selective loss of *Vmat2* transcripts in the midbrain (D, left) but not raphe nucleus (D, right) in *DAT-iCreER x Vmat2^fl/fl^*mice compared to controls (C). Scale bars, 250 μm (VTA), and 100 μm (Raphe). (E, F) Adult *DAT-iCreER* and *DAT-CreER x Vmat2^fl/fl^*mice were tamoxifen-injected daily for 5 days starting on day 4 of the running wheel recording to produce ^DAT^Vmat2KO and respective control animals. Traces along-side the actograms show the scaled-average continuous wavelet transform spectrum in the circadian (20-27 hr) and infradian (27-96 hr) range. ^DAT^Vmat2KO mice (F) but not controls (E) showed a gradual reduction in locomotor activity. Addition of Meth (indicated by orange background hues) to the drinking water rescued activity and led to the emergence of infradian components in both controls and ^DAT^Vmat2KO animals as evidenced by the increased spectral power in the infradian range (solid trace).

**Figure S5.**
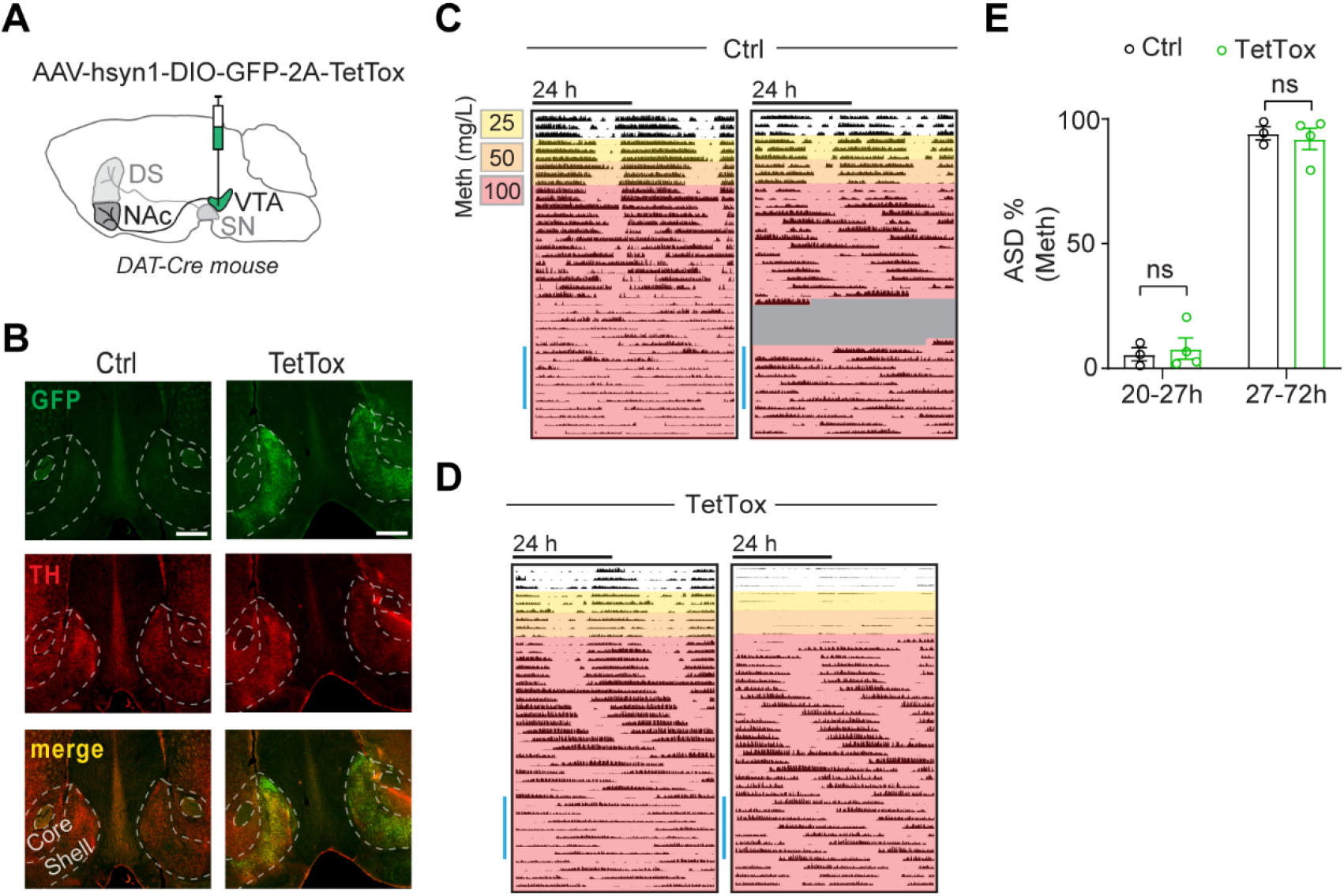
Tetanus toxin-mediated silencing of ^VTA^DA neurons does not abrogate the 2ndC induction capacity. Related to Figure 5. (A) AAV-hSYN1-DIO-GFP-2A-TetTox was bilaterally injected into the VTA of *DAT-Cre* mice. (B) Immuno-labeling for TH and GFP demonstrates effective transduction of NAc**-**shell projecting TH+ fibers. Scale bars, 500 μm. (C, D) Representative actograms of (C) Ctrl (top) vs (D) ^VTA^TetTox (bottom) mice showing running wheel activity in response to Meth-supplemented drinking water indicated by color shading. Greyed area, data loss. (E) ASD analysis of periodograms derived from the time span indicated by blue bar next to actograms. Infradian rhythmicity under Meth was indifferent between ^VTA^TetTox mice and controls, suggesting that ^VTA^DA neuronal silencing does not abrogate the 2ndC emergence capacity. Mean ± SEM. n= 3-4. Two-way ANOVA with Bonferroni’s multiple comparison. ns, not significant.

**Figure S6.**
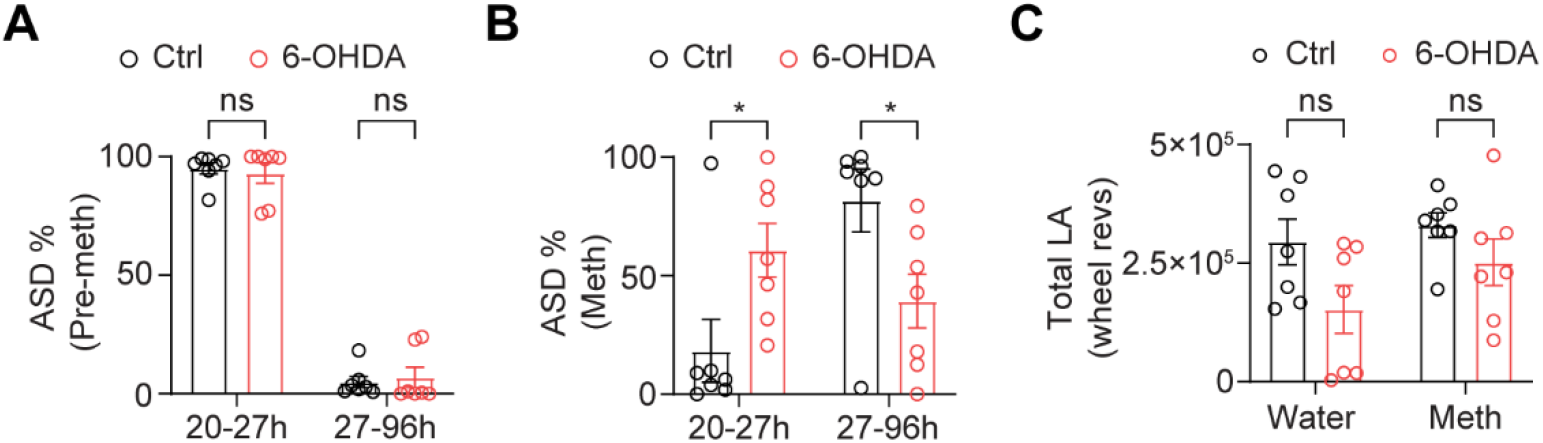
Loss of infradian rhythm generation capacity upon ablation of DA projections in the NAc. Related to Figure 7. (A, B) Percent of significant rhythmic periodicities in the circadian (20-27 hr) and infradian (27-96 hr) period range before (A) and during the final 2 weeks of Meth-exposure (B). ASD, amplitude spectral density. (C) Total locomotor activity (wheel revolutions) of 6-OHDA and Ctrl mice prior to (Water) and during Meth treatment (Meth). Mean ±SEM. n= 7. Two-way ANOVA with Bonferroni’s multiple comparison. ns, not significant; \**P*<0.05.

